# Improved model representation of the photosynthetic light reactions reduces estimates of global gross primary productivity

**DOI:** 10.64898/2026.05.08.723728

**Authors:** Julien Lamour, Jérôme Chave, Jennifer Johnson, Joseph Berry, Kenneth J. Davidson, Kim S. Ely, Liang Fang, Charles D. Koven, Jessica F. Needham, Ülo Niinemets, Raphaël Perez, Stephanie C. Schmiege, Zhihong Sun, Danielle A. Way, Alistair Rogers

## Abstract

The assimilation of carbon dioxide by plants can be predicted by the Farquhar, von Caemmerer and Berry model of photosynthesis. This largely mechanistic model is central to understanding how plants influence Earth’s climate. However, it represents the use of light by photosynthesis using an empirical formulation. Johnson and Berry proposed an alternative mechanistic formulation based on the functioning of the cytochrome b6f complex that includes key steps in light harvesting and electron transport. We compared both formulations using photosynthetic light response measurements from 146 C3 species spanning arctic to tropical biomes and implemented them in the terrestrial biosphere model ELM-FATES to simulate global photosynthesis. The Johnson and Berry formulation better fitted the measured response of leaf-level photosynthesis to light, and predicted lower photosynthetic rates at intermediate light levels, which decreased global estimations of terrestrial photosynthesis by 8%. Our findings support adopting the Johnson and Berry formulation to improve model representation of global carbon cycle modeling.

## Introduction

Terrestrial biosphere models (TBMs) represent the exchange of carbon dioxide and water vapor between plants and the atmosphere by simulating the gas exchange of individual leaves aggregated over entire ecosystems. At the heart of these models, photosynthesis is represented by the Farquhar, von Caemmerer and Berry equations (*1*) and subsequent developments (*2*) (hereafter, FvCB model). These equations model the effect of the environment, notably light, CO_2_ concentration, and temperature, on the CO_2_ assimilated by leaves. Estimates of the gross primary productivity (GPP) are highly dependent upon the assumptions and mechanistic foundations underlying model representation of leaf-level photosynthesis. Therefore, these models are crucial for accurately simulating ecosystem and global fluxes.

Photosynthesis is commonly described as consisting of two sets of reactions, the light reactions and the dark, or light-independent reactions. The light reactions convert incident light to chemical energy through electron transport (*J*). This process, regulated by the cytochrome b_6_f complex (*3*), provides the energy and reductant necessary for the dark reactions, which assimilate CO_2_ into sugars. The dark reactions are regulated by the enzyme rubisco. Within the FvCB model, the equations describing the dark reactions are derived from the mechanistic understanding of the kinetic properties of rubisco. In contrast, the light reactions are described empirically.

Over the last fifty years, versions of Equation (1) have become the most widely used empirical representation of the response of *J* to irradiance (*Q*) (*2, 4*). At low irradiance, *J* is proportional to the absorbed irradiance (*αϕQ*), where α is the leaf absorptance (the fraction of incident irradiance absorbed by the leaf), and □ is the maximum quantum yield of the absorbed light (mol of electrons per mol of absorbed photons). At high irradiance, electron transport approaches an asymptote, *J*_max,_ the maximum potential electron transport rate (where *J* = *J*_max_, µmol electrons m^-2^ s^-1^).

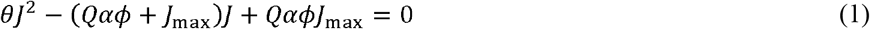

The transition between the initial slope (*J =*α□*Q*) and the asymptote (*J* = *J*_max_) of the light response is described by *θ*, a dimensionless curvature factor. A value of *θ* = 1 represents an abrupt transition, whereas lower values of *θ* represent smoother transitions (Fig. 1). The special case where *θ* = 0 corresponds to a rectangular hyperbola (*5*) and negative values are also possible (Fig. 1). While Equation (1) describes the shape of the light response, it does not explain how it emerges from the underlying biophysical and biochemical factors that control photosynthesis under light-limited conditions. In this way, Equation (1) is fundamentally different from the other equations in the FvCB model that explain both the shape of the observed responses and how it emerges from the biochemical properties of the enzyme rubisco.

**Fig. 1.**
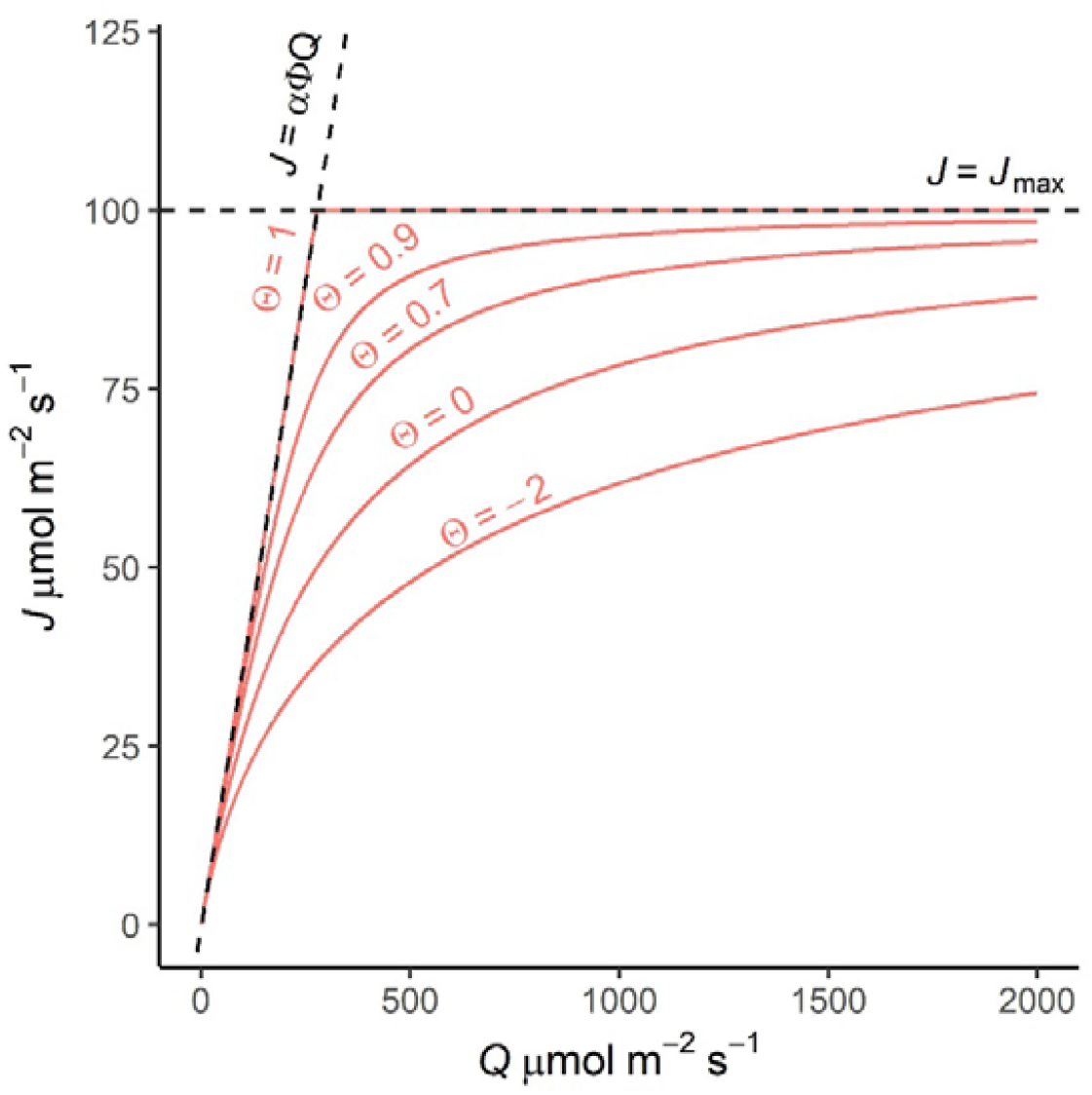
Effect of the empirical curvature parameter (θ) on the response of electron transport rate (*J*) to irradiance (*Q*). The horizontal dashed line corresponds to the maximum potential rate of electron transport (*J*_max_), the limit of *J* at infinite irradiance. The sloping dashed line corresponds to a regime where *J* is proportional to the absorbed irradiance (*αϕQ*), where α is the leaf absorptance, and □ is the maximum quantum yield of absorbed light. The *J* response curves in red correspond to empirically smoothed curves where *θ*, the curvature factor, smooths the transition between the initial slope (*J = αϕQ*) and the asymptote (*J*_max_). A value of one corresponds to an abrupt transition (no smoothing), whereas lower values, including negative values, represent smoother transitions.

In most TBMs and ecophysiological applications, *θ* is conceptualized as a universal parameter that is fixed at a constant value (Supplementary Text, table S1). In fact, this parameter is rarely estimated (*6, 7*). Instead, parameterization of the FvCB model with empirical measurements is typically accomplished by measuring the response of photosynthesis (*A*) to intercellular CO_2_ concentration (*C*_i_), commonly known as an *A*-*C*_i_ curve (*8, 9*). Photosynthesis can be limited by the carboxylation rate of rubisco (*A*_c_), the electron transport rate (*A*_j_), or the use of the triose phosphate end product (*A*_p_). The region of the *A-C*_i_ curve where electron transport limits the photosynthetic rate (*A*_j_) is used to estimate *J* (*10*), and *J*_max_ is then extrapolated using Equation (1), typically with *θ* fixed at 0.7. However, while *A*-*C*_i_ curves are useful for estimating *J*_max_, they cannot be used to estimate *θ*. For this purpose, light response curves of photosynthesis (*A*-*Q* curves) with sufficient light levels in the intermediate light range are needed. Unfortunately, these measurements are seldom made. This measurement gap and the empirical nature of *J* mean that the prediction of the rate of light-dependent reactions of the FvCB model is more empirical, less tested, and represents a potential weakness for the predictions of ecosystem-scale CO_2_ fluxes.

Recently, to place analysis and prediction of the light response on a stronger foundation, Johnson and Berry (*11*) described a model (hereafter, JB model) where the model representation of *J* is based explicitly on the operation of the cytochrome b_6_f complex, which mediates electron transport between photosystem II (PSII) and photosystem I (PSI). In this new formulation, the dependence of electron transport on irradiance is described by Equation (2),

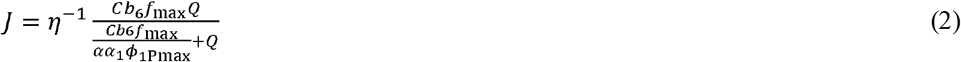

where *Cb*_6_*f*_*max*_ (µmol electron m^-2^ s^-1^) is the maximum rate of electron flow through the cytochrome b_6_f complex, *α*_1_ is the fraction of absorbed irradiance absorbed by PSI, ϕ_lPmax_ is the maximum photochemical yield of PSI (mol mol^-1^) and *η* (mol mol^-1^) quantifies the ratio of PSI to PSII electron transport rate, which depends on the leaf chloroplastic CO_2_ concentration (*C*_c_, µmol mol^-1^). The term η is expressed as a function of the coupling efficiency of linear electron flow (*n*_L_=0.75 mol ATP mol^-1^ electrons) and the coupling efficiency of cyclic electron flow (*n*_c_ =1 mol ATP mol^-1^ electrons) as follows,

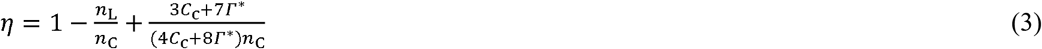

where *Γ*\* (µmol mol^-1^) is the CO_2_ compensation point.

The JB model has several advantages over the FvCB model. Its improved biochemical foundation avoids the need for the empirical parameter *θ*. Furthermore, the JB model was designed to represent leaf radiative fluxes associated with photosynthesis and particularly the simulation of chlorophyll fluorescence. Since chlorophyll fluorescence can be measured at scales ranging from the leaf to the landscape (*9, 12–14*), the JB model could provide a pathway to link fluorescence measurements made at many different scales to photosynthetic CO_2_ assimilation. Therefore, incorporating the JB formulation of *J* in the representation of photosynthesis presents an opportunity to evaluate, constrain, or calibrate vegetation models using solar-induced fluorescence measurements (*15, 16*). However, replacing the FvCB model formulation with the new formulation from Johnson and Berry in TBMs poses several challenges. First, there currently is a lack of broad empirical evidence that the JB model improves our ability to represent the response of photosynthesis to light. Second, to enable implementation of the JB formulation in TBMs, it is necessary to understand how the JB model’s parameters vary across species and environments and might relate to existing understanding of analogous parameters in the FvCB model. Finally, in the FvCB model, *J* only depends on irradiance, but in the JB formulation, *J* also depends on *C*_c_ (Equation (3)). This implies a need for adjusted sequencing of calculations that describe the transport of CO_2_ between the atmosphere and the chloroplast, regulated by stomatal and mesophyll conductance.

This study aims to, (i) test whether the JB model improves the representation of photosynthetic response to irradiance using a large dataset of photosynthetic light response measurements, (ii) develop a method to relate JB model parameters to existing FvCB model parameters, facilitating implementation in TBMs, and (iii) evaluate the impact of implementing the JB formulation on the simulation of terrestrial photosynthesis using the Energy Exascale Earth System Model Land Model - Functionally Assembled Terrestrial Ecosystem Simulator (ELM-FATES) TBM (*17*).

## Results

### The JB model predicts a more gradual response of electron transport to light than the FvCB model

Here, we demonstrate that the JB equation is mathematically equivalent to the FvCB equation when *η* in JB is set to 1 and the curvature coefficient *θ* in FvCB is set to zero. When *θ* = 0 (Equation (1)), the FvCB equation simplifies to a rectangular hyperbola

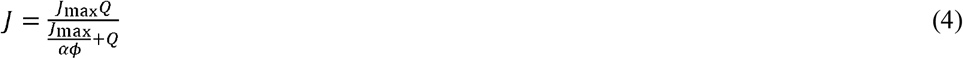

which is equivalent to the Equation (2) in the JB model for *J*_max_ *= Cb*_6_*f*_max_*η*^*-*l^ and *ϕ* = *η*^*-*l^*α*_l_*ϕ*_1Pmax_.

Although *η* in JB depends on *C*_c,_ its extreme values—where *C*_c_ is infinite and zero—range from 1 to 1.125. Thus, the theoretical range of variation of *η* with *C*_c_ is small and close to 1 (fig. S1). This range would become even smaller when considering the potential *C*_c_ encountered *in vivo*. To further test if small deviations of *η* from 1 impacted the prediction that *θ =* 0, we coupled the JB model to a stomatal model (*18*) to simulate *A-Q* curves with plausible values of *C*_c_ (assuming infinite mesophyll conductance, as is done in most TBMs). We then fit the FvCB model to the synthetic data generated by the JB model. The resulting *θ* ranged between -0.1 and -0.01, remaining very close to zero. This further demonstrated that the JB model predicts a more gradual response of electron transport rate to irradiance than the FvCB model, as currently parameterized in most TBMs, i.e., where *θ =* 0.7.

### The JB model better represents the observed response of photosynthesis to light

We compared the ability of the FvCB model and the JB model to simulate leaf-level photosynthetic response to light. We gathered 1422 *A*-*Q* curves (table S2) collected in 146 C3 species mapping to six plant functional types (*19*) spanning arctic to tropical biomes and fitted the FvCB and JB models. To ensure a fair comparison, the JB and FvCB had the same number of fitted parameters: *Cb*_6_*f*_max_ or *J*_max_, leaf day respiration (*R*_day_), and the maximum carboxylation rate of rubisco (*V*_cmax_). For the FvCB model, *θ* was fixed at 0.7. We harmonized the initial response of *J* to *Q* in the FvCB and JB models by fixing the *α*_l_ parameter of the JB model to enforce equivalence and to avoid confounding parameterization differences. Comparing their fitting errors (ΔRMSE), the JB model achieved a lower RMSE in 83% of the curves (Fig. 2A) and improved the fit quality for all six plant functional types considered in this study (Fig. 2B). The JB model also demonstrated better performance compared to the FvCB model parameterized with *θ* = 0.9 (fig. S2), an alternative parameterization used in some TBMs. Finally, to validate the theoretical equivalence of the FvCB model, parameterized with *θ* = 0, and the JB model, we compared their RMSE. Both models produced similar fits (R^2^ >0.999, slope = 1.0, intercept = 0.0, fig. S3), confirming that the FvCB model produced similar fits to the JB model when *θ* = 0. These findings confirmed that the low apparent curvature predicted by the JB model formulation was consistent with observations.

**Fig. 2.**
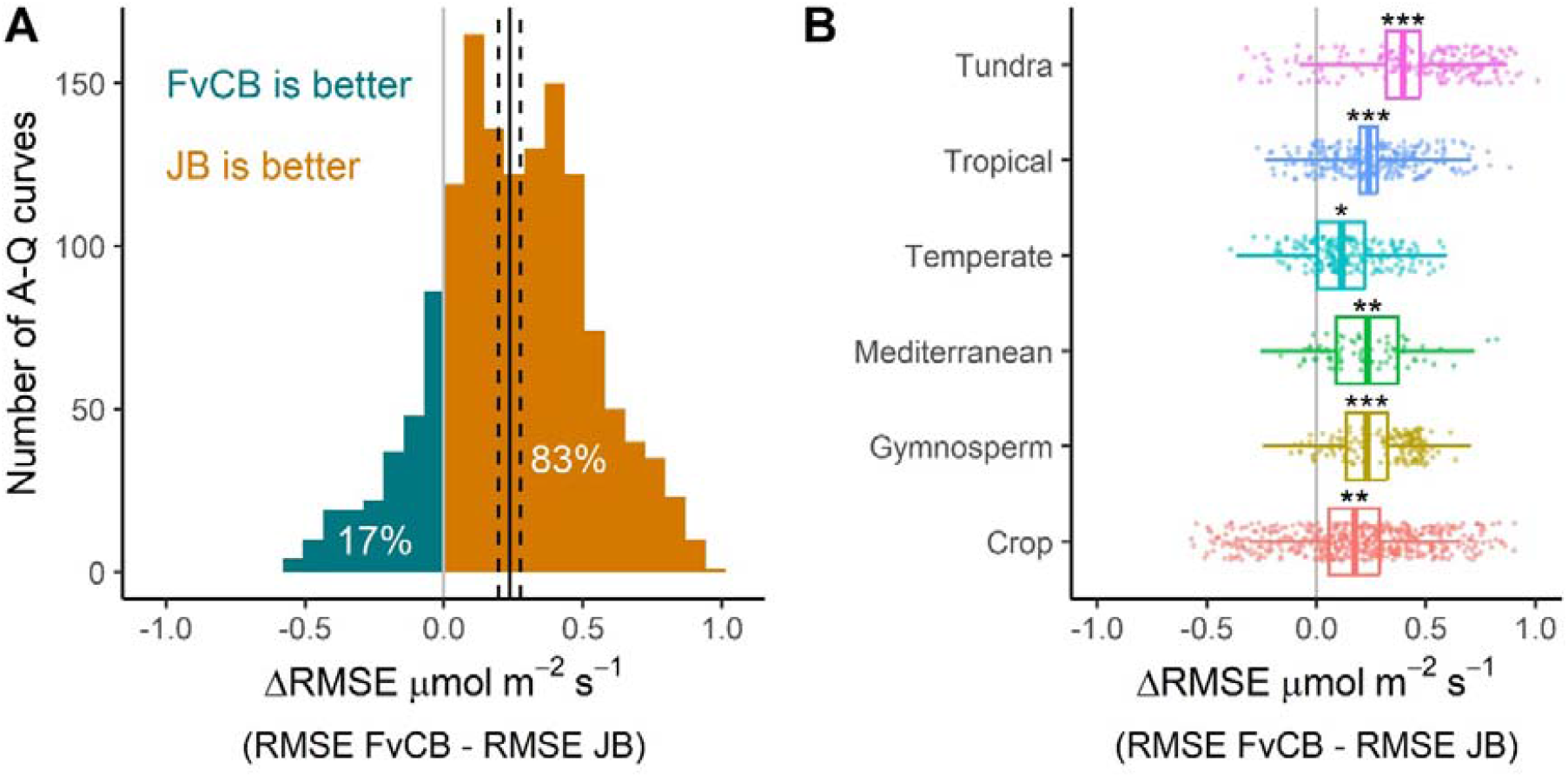
Relative performance of the JB and FvCB models for representing the response of photosynthesis to irradiance (*A-Q* curves) in C3 species. The ΔRMSE represents the difference between the Root Mean Square Error of the FvCB and JB models, where a positive ΔRMSE indicates a lower error for the JB model and vice versa. **(A)** ΔRMSE distribution in the 1422 *A-Q* curves used in this analysis. The vertical black line represents the mean ΔRMSE calculated by a mixed model that considered the species and plant functional type random effects, and the dashed lines represent the standard error of the mixed model. **(B)** Relative performance of the FvCB and JB models for each plant functional type. The top and bottom of the boxes represent the 95% confidence interval and the line is the mean estimated by a mixed model with the species represented as a random effect. The whiskers show the 95% prediction interval. The asterisks show significant difference from 0 (* P<0.05, ** P<0.01, and *** P<0.001).

### Building bridges between parameters in the FvCB and JB photosynthesis models

A challenge for implementing the JB model in TBMs is the need to explore how model parameters vary across species and environments (*20–22*). For this purpose, we proposed to relate the JB model parameters to existing FvCB model parameters so that the extensive knowledge on the FvCB parameter variation already implemented in TBMs can be used for parameterizing the JB model. For scaling the parameters, clear constraints needed to be defined to be able to establish correspondence in the free parameters of each model. We based these constraints on the conventions and experimental protocols used to estimate photosynthetic parameters. For instance, the quantum yield — corresponding to the initial slope of a light response curve — is considered a constant in most TBMs. The JB model can be parameterized to predict the same quantum yield as the FvCB model (*ϕ = η*^*–*l^*α*_l_*ϕ*_lPmax_), thereby simplifying the conversion of parameterization for the JB formulation.

For harmonizing *Cb*_*6*_*f*_max_ and *J*_max_, we have proposed a scaling based on the convention that *J*_max_ is parameterized using *A*-*C*_i_ curves and that *Cb*_6_f_max_ will be parameterized using the same protocol. This led to the constraint that both photosynthesis models necessarily predict similar *A*_j_ values and thus *J* values in the region of the *A*-*C*_i_ curve where electron transport limits photosynthesis. Therefore, by equating the *J* equations of both models, we expressed *Cb*_*6*_*f*_max_ as a function of *J*_max_, *Q, C*_c_, and all the other parameters of the *J* equations (see Methods, Equation (11)). While *C*_c_ varies across an *A*-*C*_i_ curve, its impact on *J* is negligible (*η* ≃ 1, Equation (2)), so we fixed it at a predefined value (Supplementary Text, fig. S4).

We have evaluated the scaling equation between *Cb*_6_f_max_ and *J*_max_ using 601 *A-C*_i_ curves obtained from previously published work (table S1). We found that *Cb*_*6*_*f*_max_ was 1 – 1.8 × *J*_max_ (Fig. 3A), highlighting the marked difference in the estimation of the maximum electron transport rate by the two models. Equation (11) accurately predicted *Cb*_*6*_*f*_max_ (R^2^ > 0.99, RMSE = 2.3 µmol m^-2^ s^-1^, Fig. 3B). This demonstrates that the JB model parameter *Cb*_*6*_*f*_max_ can be reliably estimated from the FvCB model parameter *J*_max_, provided the irradiance used during the *A*-*C*_i_ curves is known. Using a default value for the irradiance (e.g., 1800 µmol m^-2^ s^-1^ in Fig. 3C) still yielded good agreement (R^2^ = 0.98, RMSE = 12.3 µmol m^-2^ s^-1^, Fig. 3B). However, the estimation of *Cb*_*6*_*f*_max_ was biased when the actual irradiance at which the *A*-*C*_i_ curves were measured was markedly different than the default value.

**Fig. 3.**
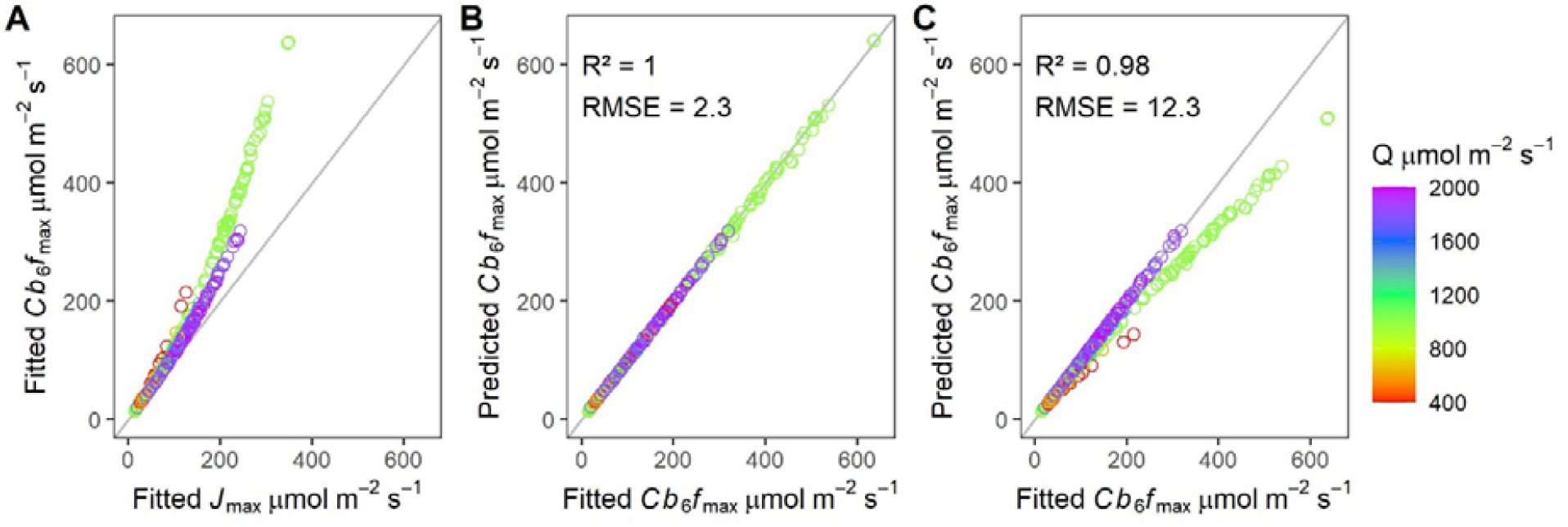
Relationship between the maximum activity of the cytochrome b_6_f complex (*V*_qmax_) and the maximum electron transport rate (*J*_max_) estimated from the response of photosynthesis intercellular CO_2_ concentration (*A-C*_i_ curves). (A)Relationship between *Cb*_*6*_*f*_max_ and *J*_max_ fitted from 601 *A*-*C*_i_ curves. **(B)** Relationship between fitted *Cb*_*6*_*f*_max_ and predicted *Cb*_*6*_*f*_max_ using the theoretical standardizing equation (Equation 11) that links *J*_max_ and *Cb*_*6*_*f*_max_, parameterized with the irradiance used during measurement of the *A*-*C*_i_ response and the default value for *C*_i_ (800 ppm). **(C)** The same relationship as in panel (B) with default values of irradiance and *C*_i_ (1800 µmol m^-2^ s^-1^ and 800 ppm, respectively). The color ramp gives the irradiance *Q* (µmol m^-2^ s^-1^) at which the *A*-*C*_i_ curves were measured.

### Effect of the models on simulations of leaf-to-terrestrial photosynthesis

As demonstrated earlier, the JB model exhibits a lower apparent curvature parameter (*θ* ≃ 0) than that which is currently implemented in most vegetation models that employ the FvCB formulation (0.7 or 0.9). To evaluate the implications of this difference for photosynthesis simulations, we harmonized the initial response of *J* to *Q* in the FvCB and JB models by fixing the *α*_l_ parameter of the JB model. We also scaled *Cb*_*6*_*f*_max_ to an equivalent *J*_max_ consistent with FvCB parameterization conventions, ensuring identical photosynthetic rates at high light (as derived from *A*-*C*_i_ curves). Simulations resulted in a more gradual photosynthesis light response for the JB model, reflecting the lower apparent curvature of simulated electron transport rate (Fig. 4). Consequently, the JB model projected a lower leaf-level photosynthetic rate at intermediate irradiance levels (Fig. 4) under photosynthetic light limitation.

**Fig. 4.**
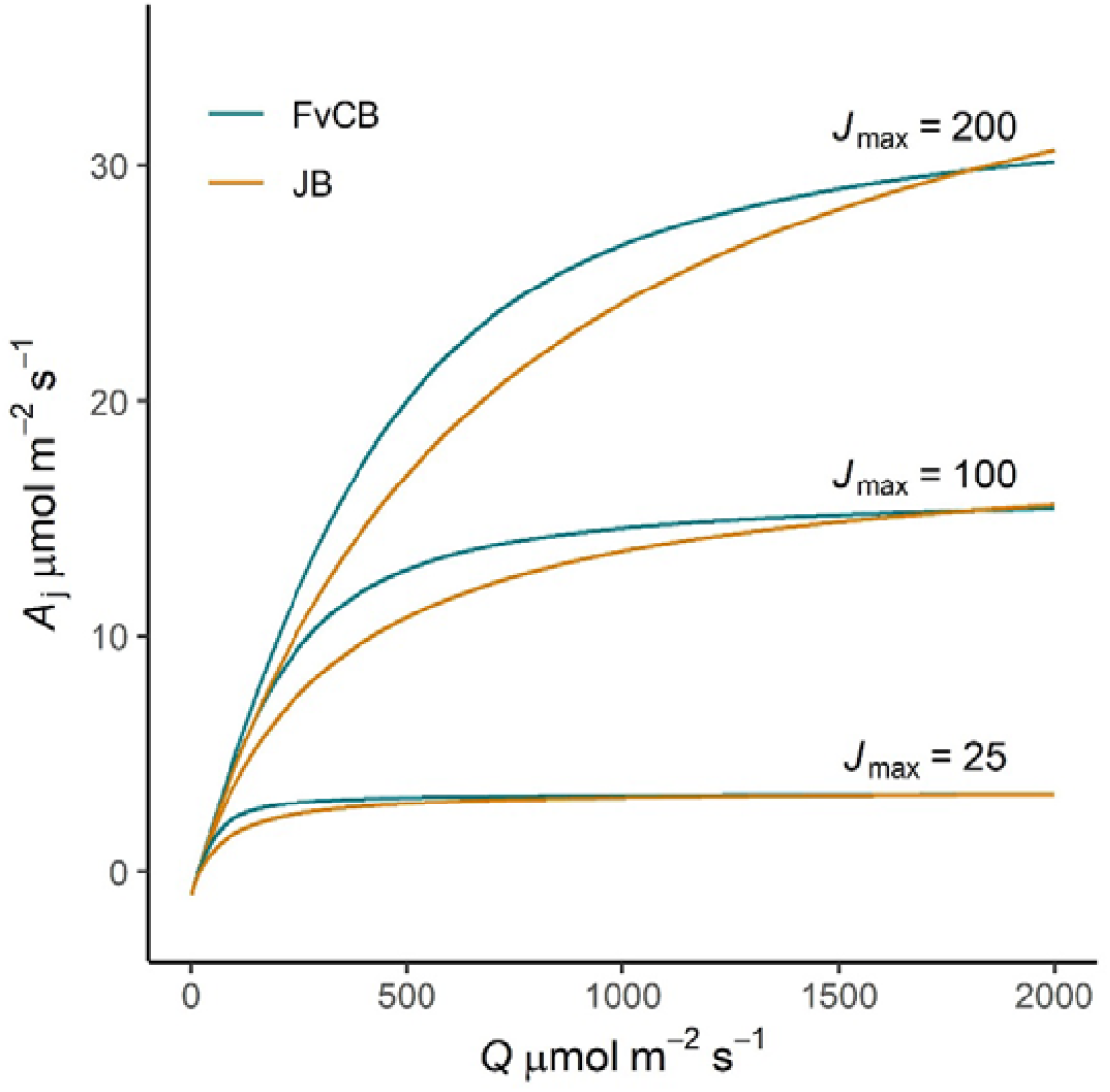
Effect of the photosynthesis model on the electron transport limited photosynthesis (*A*_j_) response to irradiance (*Q*). Blue lines correspond to the Farquhar, von Caemmerer, and Berry (FvCB) model, parameterized with *θ* = 0.7 and three different rates of maximum potential electron transport (*J*_max_, µmol m^-2^ s^-1^). Orange lines represent the Johnson and Berry (JB) model. The JB model parameters were standardized to match the slope at the origin of *A*_j_ versus *Q* to that simulated with the FvCB model. The maximum activity of the cytochrome b_6_f complex (*Cb*_*6*_*f*_max_) of the JB model was set for enforcing equivalence of *A*_j_ at 1800 µmol m^-2^ s^-1^.

We used the ELM-FATES model to further investigate the impact of model choice on the simulation of terrestrial photosynthesis. Global simulations were achieved by running ELM-FATES in a mode of operation where vegetation structural properties were prescribed by a dynamic dataset of global satellite observations (*23*). This was necessary to identify the direct effects of modifying the electron transport formulation on GPP without feedbacks between GPP and vegetation structure. Furthermore, the JB model parameters were varied by plant functional type based on their scaling to the FvCB parameters. The annual global GPP for ELM-FATES with the default FvCB formulation was 115.1 Pg C yr^-1^ (Fig. 5A). When the JB model formulation was implemented and used in place of the FvCB formulation (Fig. 5B), GPP was 8% lower (GPP = 106.0 Pg C yr^-1^ for the JB model). Differences were the strongest in the tropics, reflecting the higher GPP in this part of the world (Fig. 5C).

**Fig. 5.**
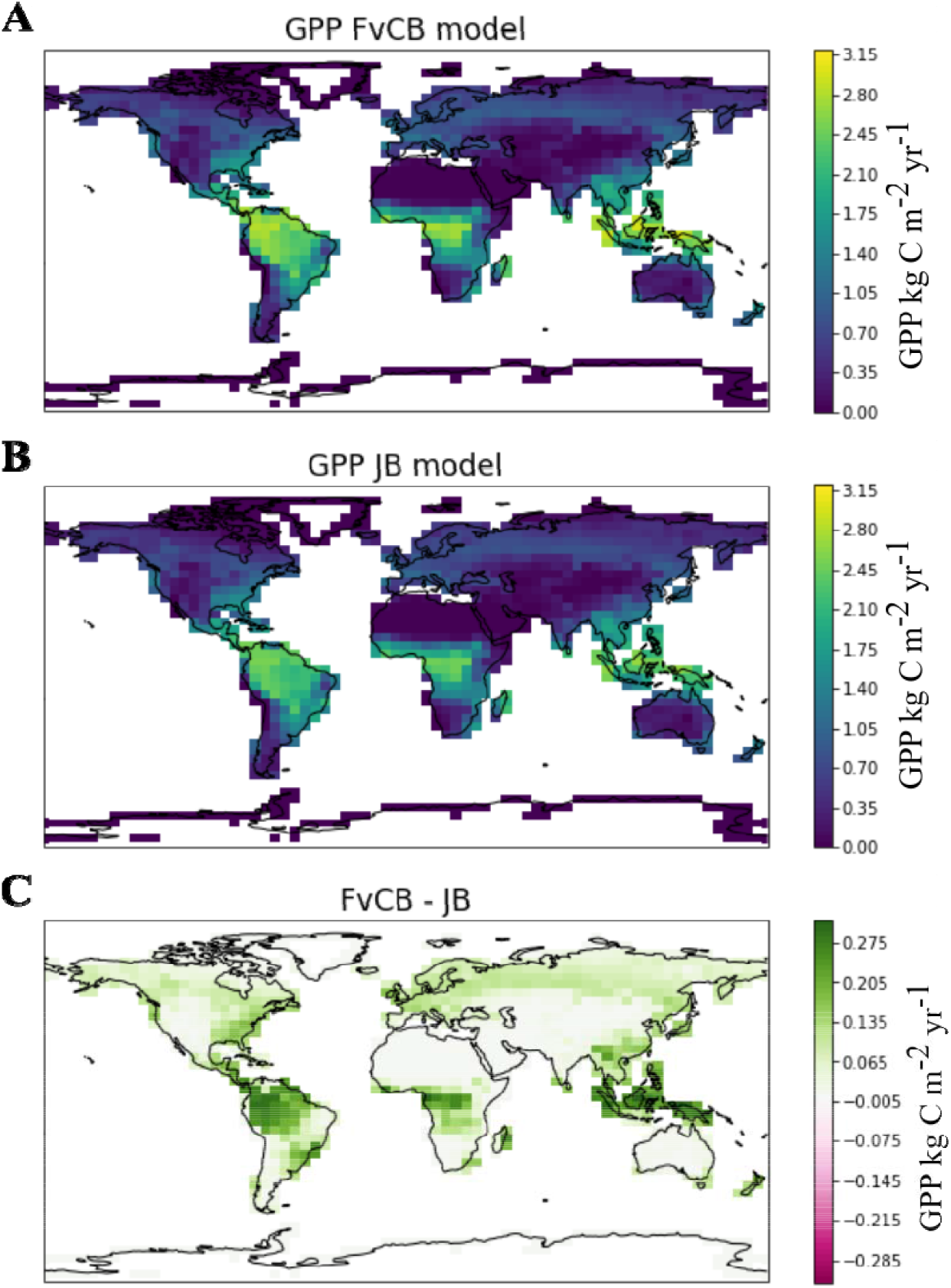
Effect of the photosynthesis model choice on global annual terrestrial gross primary productivity (GPP). Panels show model output from the terrestrial biosphere model ELM-run in a reduced complexity mode to isolate the effects of changes in model representation of photosynthesis. (**A)** GPP for the default model with the Farquhar, von Caemmerer and Berry (1980) formulation for electron transport. **(B)** GPP simulated with the newly implemented formulation for electron transport presented by Johnson and Berry (2021). **(C)** Difference in GPP between the two approaches.

## Discussion

The JB model represents a theoretical improvement upon the FvCB model by explicitly integrating the role of cytochrome b_6_f in electron transport, offering a more mechanistic description of the light reactions. We have shown that the JB model predicts a more gradual photosynthetic light response than is currently modeled in most TBMs, and this was supported by a multi-biome evaluation of the response of leaf-level photosynthesis to light. Simulations in a global model demonstrated that the implementation of the JB model formulation lowered terrestrial GPP predictions by 8% compared to the usual representation. Furthermore, we have shown that the technical challenges of implementing the JB model formulation in TBMs can be addressed. In particular, the existing extensive parameterization of leaf photosynthetic parameters can be readily adapted for the JB model. Given the critical importance of accurately modeling the effect of light on ecosystem productivity (*24, 25*), the present work will hopefully pave the way toward the implementation of the JB model formulation in leaf-level, whole-plant, and global models of photosynthesis.

Our results support the simulation of a more gradual light response by the JB model than the response currently modeled in most TBMs that use the FvCB model with *θ* = 0.7 to 0.9. Given the widespread use of the FvCB equations for studying photosynthesis in ecophysiology, it may be surprising that this fundamental mismatch has not received much attention. We argue that this oversight stems from two key challenges: (1) the scarcity of *A-Q* measurements where *A*-*C*_i_ curve is the most widely adopted protocol for parameterizing the FvCB model, and (2) the technical and conceptual difficulties in estimating *θ*.

Estimating *θ* requires analysis of *A*-*Q* curves, but its derivation is not straightforward. This is because *J*, defined by Equation (1) or Equation (2), is a potential rate that is only realized if the electron transport rate limits photosynthesis (*1*). *In vivo*, at high irradiance, the actual electron transport rate is typically below the maximum potential *J*_max_ predicted by Equation (1) or (2). Instead, when *A* is light-saturated at ambient CO_2_ concentration, photosynthesis and electron transport rates are usually limited by the maximum carboxylation rate of rubisco (*V*_cmax_) (*11, 26*), i.e., *A* is limited by *A*_c_. Indeed, this observation enabled the development of a simplified technique — the one-point method — for measuring *V*_cmax_ (*27–29*). The prevalence of *A*_c_ limited *A* at high irradiance is also consistent with a coordinated allocation of resources in the leaf for the electron transport chain, and for carbon assimilation, which is used to predict *V*_cmax_ in some vegetation models (*30, 31*) (Coordination theory (*32*)). Crucially, if photosynthesis is limited by the carboxylation rate at high irradiance, this implies that in *A*-*Q* curves, the photosynthetic rate is limited by the electron transport rate at low irradiance (*A*_j_) and by the carboxylation rate of rubisco (*A*_c_) at high irradiance, i.e., the observed *J* is below *J*_max_. Therefore, if we can’t observe the true *J*_max_ *in vivo*, how can we also estimate the nature of the transition from *J =*α□*Q* to *J*_max_? Indeed, the elusive nature of *J*_max_ and *θ* combined with the dominance of *A*_c_ limitation at high light led Collatz et al. (*33*) to only model the light-limiting part of electron transport (*J* = α□*Q*), an assumption used in some TBMs (*34, 35*).

Further complicating matters, a wide range of *θ* values have been reported, from negative to nearly one (*7, 36–42*) using various protocols for its estimation. In a few studies, *θ* was estimated using Equation (1) (*37, 38, 40*), but many studies used the apparent curvature of the response of *A* to *Q* (*36, 41, 42*), using Equation (5), which is one of the early models of photosynthesis (*5*):

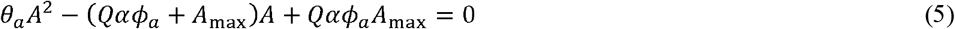

However, *θ*_a_ does not necessarily equal *θ* (Supplementary Text), and the use of Equation (1) for modeling *J* was primarily a historical choice based on the similarity with Equation (5) (*43*). The technical and conceptual challenges associated with estimating *θ*, combined with the better representation of light-response curves by the JB model, provide compelling justification for adopting the JB model in TBMs and ecophysiological applications.

A major barrier to the adoption of the JB model in large scale models is the need to understand how its parameters vary across species and environmental conditions. Our results demonstrated that the JB model parameter *Cb*_*6*_*f*_max_ can be reliably estimated from the FvCB model parameter *J*_max_. This ability to inform parameterization of the JB model will enable its initial implementation in vegetation models that currently include model representation of variation in *J*_max_ with plant functional type or environment (*21, 44, 45*). The accuracy of the *Cb*_*6*_*f*_max_ estimation relied on knowledge of the irradiance at which the *A*-*C*_i_ curves were performed. Most empirical studies used to parameterize TBMs are based on measurements of top of canopy leaves, where the measurement irradiance is typically set at 1800 µmol m^-2^ s^-1^, which enables the estimation of *Cb*_*6*_*f*_max_. An alternative for the parameterization of *Cb*_*6*_*f*_max_ in TBMs is to directly calibrate the JB model using *A*-*C*_i_ curves. This requires that *A*-*C*_i_ curves for all the species and biomes currently represented in TBMs are available. This rigorous approach for parameterizing TBMs would ensure that the same set of equations and assumptions is used to estimate parameters and predict photosynthesis, avoiding biases that result from mixing and matching equations (*21*).

The GPP simulated by ELM-FATES with the JB photosynthesis model was 8% lower than that obtained with the FvCB model. This difference arises because the JB model predicts lower photosynthetic rates at intermediate irradiance under light-limited conditions, and this effect propagates to the global scale. Note that the veracity of the absolute projected global GPP is difficult to assess, but the impact of adopting the JB formulation is clear. Indeed, efforts to constrain estimates of global GPP are ongoing and challenging (*46, 47*). The core value proposition of the JB model to Earth science applications is the potential to add an additional constraint on estimated GPP via solar-induced chlorophyll fluorescence since the JB equations also predict leaf-level radiative fluxes that can be readily implemented in TBMs. By demonstrating that the JB model accurately predicts gas exchange measurements, we have completed a first critical step necessary to unlock its full potential for advancing Earth system modeling through direct fluorescence-based assessment.

## Materials and Methods

### Model comparison and equivalence between parameters

In the FvCB model (*1, 2, 39*), the net CO_2_ assimilation rate (*A*_n_, µmol CO_2_ m^-2^ s^-1^) is determined as the minimum of three potentially limiting processes: the rubisco-limited assimilation rate (*A*_c_), the electron transport-limited assimilation rate (*A*_j_), or the triose phosphate-limited assimilation rate (*A*_p_), where *R*_day_ is the daytime respiration rate (µmol CO_2_ m^-2^ s^-1^) that is not attributable to the photorespiratory pathway:

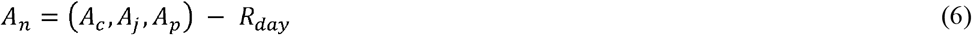

*A*_j_ is of interest here, and is described by Equation (7):

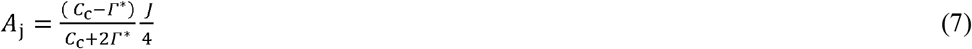

where *J* is the linear electron transport rate through Photosystem II (µmol m^-2^ s^-1^), *Γ* * is the CO_2_ compensation point in the absence of non-photorespiratory respiration in the light (µmol mol^-1^) and *C*_c_ is the chloroplastic CO_2_ concentration (µmol mol^-1^) that we assumed to be equal to the intercellular CO_2_ concentration (*C*_i_) for the purposes of this analysis (i.e., assuming mesophyll conductance is infinite).

The response of *J* to *Q* is modeled as a quadratic equation in the FvCB model (Equation (1)), whose solution is presented as Equation (8), or Equation (4) when *θ* (dimensionless) is zero.

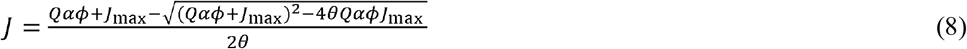

Note that since *A*_n_ is the minimum of three potential rates, *J* given by Eqn 8 or Eqn 4 is also a potential rate. The actual *J* if *A*_c_ or *A*_p_ limit photosynthesis is given by solving for *J* in Eqn 7 using *A*_c_ or *A*_p_ in place of *A*_j_.

The maximum quantum yield of absorbed light, □ (mol electron mol^-1^ photon), is considered constant for many applications and can be further described by Equation (9) (*2, 48*), where *f* is the fraction of light that is not used for photochemistry and β is the fraction of absorbed light that reaches PSII relatively to PSI.

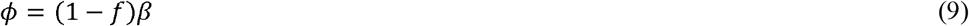

The variable □ is also the partial derivative of *J* with *αQ* at *Q* = 0, i.e., the initial slope of *J* with α*Q*.

In the JB model, *J* is computed from the properties of the cytochrome b_6_f complex (Equation (2)). The maximum quantum yield of absorbed light, as defined in the FvCB model, is described by equation (10).

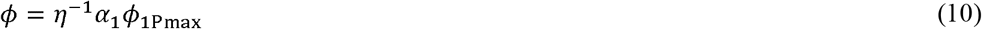

In Equation (10), *ϕ*_lPmax_ is the maximum photochemical yield of the photosystem I (0.96 mol energy dissipated mol^-1^ energy absorbed) and η is a variable quantifying the ratio of PSI to PSII electron transport rate, which depends on *C*_c_ (Equation (3)). The variable *α*_l_ (mol photon mol^-1^ absorbed *Q*) is the fraction of absorbed light that reaches PSI relatively to PSII (*α*_l_ *=*1 – *β*). Note that Johnson and Berry originally used the absolute absorptance of PSI in mol of absorbed *Q* per mol of incident *Q* at the leaf surface. We chose to use *αα*_l_ in Equations (2 and 10) to harmonize notations. Johnson and Berry also proposed equations to model dynamic cross sections of PSII and PSI that depend on light intensity. These equations were developed to represent a regime where chloroplast movements decrease light absorption. Here, we used the static case where *α*_l_ remains constant.

Equation (10) can be used to set *α*_l_ in the JB model such that the quantum yield is identical to that of the FvCB model. The other parameters, *ϕ*_lPmax_ and the ones defining *η* (Equation (3)), should be considered constant for application in vegetation models (*11*).

The asymptote of *J* is *J*_max_ in the FvCB model and *Cb*_*6*_*f*_max_*η*^*–*l^ in the JB model. It is possible to use this property to parameterize the JB model such that both models have the same asymptotic limit. However, this is not useful for most applications. In practice, *J*_max_ is extrapolated from *J* estimated from the measured response of *A* to *C*_i_ (*A*-*C*_i_ curves) in the domain where photosynthesis is *A*_*j*_ limited. The light irradiance used in the *A*-*C*_i_ curves is typically set at a high level (*Q*_sat_). Therefore, to find a relation between *Cb*_*6*_*f*_max_ and *J*_max_, a more relevant approach is to find *Cb*_*6*_*f*_max_ such that *J* estimated by the JB model is similar to *J* estimated by the FvCB model. Using this constraint leads to Equation (11) for *Cb*_*6*_*f*_max_:

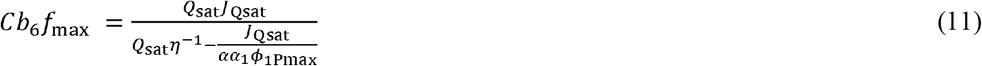

where *J*_Qsat_ is the electron transport rate estimated by the FvCB model at *Q*_sat_. (Note that *J*_Qsat_ < *J*_max_). Comparison of the response of photosynthesis to irradiance with the JB and FvCB models

We compared *A-Q* curves simulated using the FvCB and JB equations to identify the consequences of using the two different models. Simulation of *A*_j_ requires an estimate of *C*_c_ which depends on the fluxes of CO_2_ between the leaf surface and the leaf intercellular space through the stomata and leaf epidermis (assuming infinite mesophyll conductance). *J* also depends (weakly) on *C*_c_ for the JB model (Equations (2) and (3)). We therefore modeled the diffusion of CO_2_ into the intercellular space using the Unified Stomatal Optimization conductance model (*18*). This model uses two parameters, *g*_0,_ the minimum leaf conductance, and *g*_1,_ a parameter representing the sensitivity of the stomatal conductance to photosynthesis. We used a common analytical framework to couple the USO and FvCB equations that we adapted for the JB model (Supplementary Text).

e first simulated *A-Q* curves with the FvCB model coupled with the USO model. We chose four different values for *J*_max_ (25, 100, 200, and 300 μmol m^-2^ s^-1^) and eight values for *g*_1_ (1 to 8) that span the range of values measured for C3 species (*44, 49*) and we set α to 0.85, *θ* to 0.7 and *ϕ* to 0.425 (*48*) (Fig. 4: *g*_1_ = 4, *J*_max_ = 25, 100, 200). We assumed that *R*_day_ and *g*_0_ were the same for all simulations and set them to 1.5 μmol m^-2^ s^-1^ and 0.01 mol m^-2^ s^-1^, respectively. We varied *Q* between 0 and 2000 µmol m^-2^ s^-1^. The other leaf environmental variables, *T*_leaf_ and *VPD*_leaf,_ were held constant at 25 □ and 0.95 kPa, respectively. We then simulated *A-Q* curves with the JB model coupled with the USO model. We parameterized the JB model so that *ϕ* was identical to the FvCB model (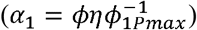). We used Equation (11) to solve for *Cb*_*6*_*f*_max_ values that would correspond to the *J*_max_ values used for the FvCB simulations, so both curves would intersect at *Q* = 1800 μmol m^-2^ s^-1^. In Equation (11), we set *Q* = 1800 μmol m^-2^ s^-1^ and *C*_i_ to the value predicted by the FvCB model for *Q* = 1800 μmol m^-2^ s^-1^. We finally fitted the *A*-*Q* curves simulated by the JB model with the FvCB model to estimate *θ* and evaluate if the departure from *η* =1 strongly impacted the apparent curvature of the JB model. Data fitting was performed using the ‘mle2’ function from the R package ‘bbmle’(*50*).

### Comparison of the FvCB and JB performance in representing photosynthetic light response

We fitted *A-Q* curves measured in 146 C3 species (table S2) and different environments against Equation (6) using Equation (1) or Equation (2) for *J*. The data were collected with commonly used measurement methods for gas exchange as described in the references cited in table S2. The irradiance was changed in several steps with sufficient time for photosynthesis to acclimate to the new environment (typically a few minutes, “rapid” light curves) in controlled conditions of temperature and CO_2_ concentration around ambient atmospheric concentration. We have also used datasets where sufficient time was given for the stomata to adapt to the new light conditions (usually more than 10 minutes, “slow” light curves). For this analysis, we only considered *A*-*Q* curves with 100 < *C*_i_ <1000 µmol mol^-1^, *g*_sw_ > 0.01 mol m^-2^ s^-1^, and relative humidity below 85 % (*8*). We removed intermediate measurements in the curves that were added to extend the time needed to reach steady-state, and we excluded curves with fewer than six points.

We fitted the data using an approach commonly used for *A*-*C*_i_ curves (*8, 9*). We considered that *A*_j_ was limiting *A*_n_ at low *Q* and that *A*_c_ could be limiting at high *Q*. We fitted the *A*-*Q* curves using *A*_j_ only and refitted the data using both *A*_c_ and *A*_j_ limitations. We selected the most parsimonious model according to the lowest Akaike information criterion (AIC). As a result, we estimated *J*_max_, *Cb*_*6*_*f*_max,_ and *R*_day_, as well as *V*_cmax_ if *A*_n_ was *A*_c_ limited at high *Q*. The fit was performed using the ‘mle2’ function from the R package ‘bbmle’(*50*). For each *A-Q* curve, the transition between light-limited and carbon-limited regimes was assigned as a result of the fitting procedure, so the set of parameters maximized the likelihood of the estimation. The root mean square error (RMSE) was calculated for each *A*-*Q* curve using both models.

We compared the FvCB and JB models by calculating the difference in their RMSE. A positive difference (ΔRMSE) indicated a lower error for the JB model and vice versa. To statistically test if ΔRMSE significantly differed from zero while accounting for the unbalanced number of *A-Q* curves per species and PFT, we used a mixed-effects model with species as a random effect nested within the PFT. We also tested if the ΔRMSE significantly differed from zero for each PFT using the PFT as a fixed effect and the species as a random effect.

For these fittings and comparison of model performance, we used standard parameters for α, *θ*, and *ϕ* with the FvCB model (0.85, 0.7, and 0.425, respectively), as well as equivalent parameters for the JB model (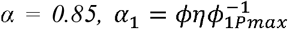, Fig. 2). Since some vegetation models use a lower value of *ϕ* (0.372) and a higher value of *θ* (0.9, Supplementary Text, table S1), we also considered this alternative parameterization to fit and compare the performance of the JB and FvCB models (fig. S2).

### Evaluation of a standardizing equation to predict *Cb*_***6***_***f***_**max**_ **from *J***_**max**_

We evaluated the performance of Equation (11) to predict *Cb*_*6*_*f*_max_ based on *J*_max_ using the fitting of *A*-*C*_i_ curves. We used 7 previously published datasets collected with commonly used *A*-*C*_i_ measurement protocols (table S2). We fitted the 601 *A*-*C*_i_ curves with the FvCB and JB models to estimate *J*_max_ and *Cb*_*6*_*f*_max_ using standard protocols (*8*). We considered that *A*_c_ was limiting at low *C*_c_ and that *A*_j_ and/or *A*_p_ could limit *A*_n_ at higher *C*_c_. The transition between *A*_c_, *A*_j_, and *A*_p_ was determined automatically by the fitting procedure. The *A*_j_ and *A*_p_ limitations were only considered if they improved the fitting of the curves according to the AIC criterion. In total, we therefore estimated up to four parameters, *R*_day_, *V*_cmax_ as well as *J*_max_ and the triose phosphate limitation if *A*_j_ and *A*_p_ were limiting at high *C*_c_. We also considered the case where *A*_c_ limits *A*_n_ without *A*_j_.

We compared the estimated *Cb*_*6*_*f*_max_ with the predicted values from Equation (11) using a standard value for *C*_i_ of 800 ppm and *Q*_sat_ set to the value used in each *A*-*C*_i_ curve. We also evaluated the performance of Equation (11) parameterized with a standard value of *Q*_sat_ of 1800 µmol m^-2^ s^-1^, a typical value used to measure *A-C*_i_ curves.

### Global terrestrial photosynthesis simulations using ELM-FATES TBM

We tested the effects of the novel JB model formulation on global GPP using the TBM ELM-FATES (*17, 24*), a cohort-based vegetation demographic model. We ran global ELM-FATES simulations in Satellite Phenology mode, in which leaf area index and vegetation height are prescribed from global datasets (*23*). This allowed us to isolate the effects of the electron transport model on gross primary productivity, without feedback from changes in vegetation structure. We ran the simulation on a 4 × 5 degree resolution grid, with climate forcing from the Global Soil Wetness Project 3 (*51*). In the implementation of the JB model in ELM-FATES, we assumed that the initial slope of *J* with α*Q* of the two models was the same (Equation (10)) and that the temperature response of *Cb*_*6*_*f*_*max*_ matched the temperature response of *J*_max_ in the FvCB model. For conversion of PFT specific *J*_max_ parameterization to Cytochrome b_6_f parameterization, we used equation (11) assumed *Q*_sat_ of 1800 μmol m^-2^ s^-1^ (typical irradiance used for *A*-*C*_i_ curves) and a leaf absorptance of 0.85.

## Author contributions

Conceptualization: JL, AR, JC

Methodology: JL, AR, JC

Investigation: JL, AR, JC, JFN

Visualization: JL, AR

Writing—original draft: JL

Writing—review & editing: All authors

## Data and materials availability

The code and data that support the findings of this study are accessible on GitHub (https://github.com/JulienLamour/Comparison_FvCB_JB).

FATES code is freely available from the FATES GitHub repository at https://github.com/NGEET/fates. All simulations were run on JFN’s fork of FATES on the jb-electron branch with commit 3923079, https://github.com/JessicaNeedham/fates/tree/jb-electron. The implementation of the JB model is now integrated into the main branch of FATES.

