## Supplementaty information for "Improved model representation of the photosynthetic light reactions reduces estimates of global gross primary productivity"

Supplementary Text

Empirical equations of electron transport rate used in terrestrial biosphere models

Several empirical relationships are used to model electron transport rate (*J*) in terrestrial biosphere models (TBMs) that are used to simulate vegetation within global climate models. Many use the nonrectangular hyperbola (Equation (S1)), but Smith’s equation in Harley *et al.* (1992) is also used (Equation (S2)). Some TBMs also use a linear equation (Equation (S3), Collatz *et al.*, 1991) that is not bounded (i.e., no *J*_max_) with the assumption that at high irradiance, the maximum carboxylation rate limits photosynthesis (*A*_c_) and so modeling a finite limit for electron transport is not necessary.

$J=\frac{Q\alpha\phi+J_{max}-\sqrt{{(Q\alpha\phi+J_{max})}^{2}-4\theta{Q\alpha\phi J}_{max}}}{2\theta}$	(S1)

$J=\frac{Q\alpha\phi}{\sqrt{1+\left( \frac{Q\alpha\phi}{J_{max}} \right)^{2}}}$	(S2)

$J=Q\alpha\phi$	(S3)

In these equations, *Q* is the incident irradiance at the leaf surface, $\alpha$ is the leaf absorptance, $\phi$ is the maximum quantum yield for absorbed light, *θ* is an empirical curvature factor of the response of *J* to *Q*, also called convexity, and *J*_max_ is the maximum rate of electron transport.

Table S1 details the parametrization of *J* equations used in some TBMs, as well as the important physiological studies on which TBMs are often based upon. The quantum yield based on absorbed irradiance (*Φ*) or the apparent quantum yield based on incident leaf irradiance (*αΦ*) is reported depending on available information.

Curvature of the response of electron transport rate and photosynthesis to irradiance

Here, we compare the curvature of the response of electron transport rate (*J*) to irradiance (*θ*, Equation (S4)) with the curvature of the response of gross photosynthetic rate (*A*) to irradiance (*θ*_a_, Equation (S5))_._

$\theta J^{2}-\left( Q\alpha\phi+J_{\max} \right)J+{Q\alpha\phi J}_{\max}=0$	(S4)

$\theta_{a}A^{2}-\left( Q\alpha\phi_{a}+A_{\max} \right)A+Q\alpha\phi_{a}A_{\max}=0$	(S5)

In these equations, $\alpha$ is the leaf absorptance, $\phi$ is the maximum quantum yield of electron transport of absorbed light (mol of electrons per mol of absorbed photons), whereas $\phi_{a}$ is the maximum quantum yield of CO_2_ assimilation for an absorbed light (mol CO_2_ per mol of absorbed photon). *J*_max_ is the asymptote of *J* (µmol electron m^-2^ s^-1^) and *A*_max_ is the asymptote of *A* (µmol CO_2_ m^-2^ s^-1^).

In the FvCB model, the gross CO_2_ assimilation rate (*A*) is determined as the minimum of two (or three) potentially limiting processes: the rubisco limited assimilation rate (*A*_c_) and the electron transport limited assimilation rate (*A*_j_):

$A=\left( A_{c},A_{j} \right)$	(S6)

*A*_c_ is given by

$A_{c}=\frac{\left( C_{i}-\Gamma^{*} \right) V_{\mathrm{cmax}}}{C_{i}+K_{c}\left( 1+\frac{O_{2}}{K_{o}} \right)}$	(S7)

where $K_{c}$ and $K_{o}$ (µmol mol^-1^) are the Michaelis−Menten coefficients of rubisco activity for CO_2_ and O_2_, respectively, *C*_i_ is the intercellular CO_2_ concentration (µmol mol^-1^), and $\Gamma^{*}$ is the CO_2_ compensation point (µmol mol^-1^).

*A*_j_ is given by Equation (S8)

$A_{j}=\frac{\left( C_{i}-\Gamma^{*} \right)}{4C_{i}+8\Gamma^{*}}J$	(S8)

Equation (S5) can be obtained from Equation (S4) using Equations (S8) and (S9), and in this case, $\theta_{a}= \theta$.

$\phi_{a}= \frac{\left( C_{i}-\Gamma^{*} \right)}{4C_{i}+8\Gamma^{*}}\phi$	(S9)

The terms $\phi$ and $\phi_{a}$ are constants in Equations (S4) and (S5). For $\theta_{a}= \theta$, *C*_i_ must be constant in Equation (S9), and *A*_j_ must limit *A* over the range of irradiance. However, it is commonly accepted that *A*_j_ limits *A* at low light, but that the maximum carboxylation rate of rubisco (*A*_c_) limits *A* at high light (*1*, *26*, *27*, *48*). It is also common that *C*_i_ varies during light curves. As a consequence, $\theta_{a}$ is not a robust estimator of $\theta$, and the relation between $\theta_{a}$ and $\theta$ is impacted by the transition from *A*_j_ to *A*_c_, and the variation of *C*_i_ with *Q* during measurement of photosynthetic light curves.

Effect of the FvCB and JB models on the photosynthesis response to CO_2_

The usual method to estimate the photosynthetic parameters of the FvCB model, including *J*_max_, is to measure the response of photosynthesis to intercellular CO_2_ concentration, commonly called *A*-*C*_i_ curves, at a saturating irradiance, and to fit the FvCB model to the measurements (*9*, *53*). At low *C*_i_, *A* is limited by the carboxylation rate of rubisco (*A*_c_, Equation S7). At higher *C*_i_, the electron transport rate (*J*) limits photosynthesis (*A*_j_, Equation S8)*.* The calculation of *A*_c_ is not changed by the JB model, the only impact is the calculation of *A*_j_. Therefore, the only effect of using the JB Equations is to modify the *A*_j_ limited part of the *A*-*C*_i_ curves. We also remark that if *J* in the JB model did not depend on *C*_i_, using the JB or FvCB model would have no impact on simulated *A*_j_ (provided that the JB model is parameterized so that *J* is identical to the FvCB model). We have shown that *J* depends on *C*_i_ in the JB model because of *η,* but this effect is expected to be small (see Equations (2) and (3)). Here, we performed simulations of *A*-*C*_i_ curves with the FvCB and JB models to test this assumption.

In the simulations, we set *Q* to 2000 μmol m^-2^ s^-1^, *T*_leaf_ to 25℃, and varied *C*_i_ between 40 and 1400 ppm. We fixed *R*_day_ to 1.5 μmol m^-2^ s^-1^, and chose *V*_cmax_ values that spanned the range of values measured for C3 species at 25°C (10, 50, 120, and 200 μmol m^-2^ s^-1^). We scaled *J*_max_ based on *V*_cmax_ using three J_max_ : *V*_cmax_ ratios of 1.3, 1.67, and 2 to encompass a wide range of possible values for *J*_max_. For the JB simulations, we parameterized the JB model α_1_ such as $\phi$ was identical to the FvCB model using Equation (10). We harmonized *Cb*_6_*f*_max_ with *J*_max_ using Equation (11), where *C*_i_ = 800 ppm and *Q* = 2000 μmol m^-2^ s^-1^ represent conditions when *A*_j_ is limiting *A*.

The difference between *A*-*C*_i_ curves simulated with the FvCB and JB models was minimal (fig. S4), and the simulated curves overlapped nearly perfectly. Differences originated from *η* (fig. S1) that is close to one and nearly invariant for high values of *C*_i_ that are typically observed when *A*_j_ limits *A*. Note that in the simulations, the *A*-*C*_i_ curves simulated with the FvCB and JB models crossed at *C*_i_ = 800 µmol mol^-1^ as a result of using Equation (11) with *C*_i_ set at 800 µmol mol^-1^.

Coupling the JB photosynthesis model with a stomatal conductance model

The FvCB and JB equations that link the photosynthetic rate to irradiance (Equation (6)) depend on the CO_2_ concentration in the substomatal cavity (*C*_i_), which is regulated by the stomatal aperture. To estimate *C*_i_, it is necessary to use a stomatal conductance model. Here we use the Medlyn *et al.*, (2011) model :

$g_{\mathrm{sw}}=g_{0}+m\frac{A_{n}}{\mathrm{CO}_{2s}}$	(S10)

This model has been parametrized for numerous plant species and environmental conditions (*55*, *56*) and derives from optimization theory. Here, *g*_0_ represents a non-zero conductance at low light, *CO*_2s_ is the CO_2_ concentration at the leaf surface, and *m* depends on the stomatal parameter *g*_1_ and the leaf-to-air vapor pressure deficit (*VPD*_leaf_):

$m=1.6(1+\frac{g_{1}}{\sqrt{VPD_{\mathrm{leaf}}}})$	(S11)

Diffusion of CO_2_ from the leaf surface to the intercellular environment is described by Fick’s law of diffusion, although other formulations exist (*57*–*59*):

$C_{i}=CO_{2s}-1.6\frac{A_{n}}{g_{\mathrm{sw}}}$	(S12)

where 1.6 is the ratio of diffusivity of H_2_O and CO_2_ through the stomata.

Equations (6), (S10), and (S12) must be solved together to express *C*_i_, *A*_n_, and *g*_sw_ in relation to environmental variables. Analytical solutions to this set of coupled equations are available (*60*–*62*) for *A*_n_ expressed in the following form:

$A_{n}=\frac{\left( C_{i}-\Gamma^{*} \right) x}{C_{i}+y}-R_{\mathrm{day}}$	(S13)

where *x* and *y* take different values depending on the process limiting *A*_n_ (*A*_c_, *A*_j_, or *A*_p_). In the FvCB model, in the electron-transport limited regime, *x* is *J*/4 and *y* is $2\Gamma^{*}.$ The solutions for *C*_i_ correspond to the roots of the following quadratic Equation (S14) which can be used then to calculate *A*_n_ (Equation (S13)) and *g*_sw_ (Equation (S10)).

$ac_{i}^{2}+bc_{i}+c=0$	(S14)

where :

$a=g_{0}+\frac{m}{CO_{2s}}\left( x-R_{day} \right)$	(S15)

$b=yg_{0}+\frac{m}{CO_{2s}}(-\Gamma^{*}x-R_{day}y)-CO_{2s}g_{0}+(x-R_{day})(1.6-m)$	(S16)

$c=-yCO_{2s}g_{0}+(1.6-m)(-\Gamma^{*}x-R_{day}y)$	(S17)

This solution can’t be directly used for the JB model as *J* depends on *C*_i_. However, *A*_j_ can be written in the same form as Equation (S13) with *x* and *y* given in Equations (S18) and (S19) thus enabling the same analytical solution as for the FvCB model.

$x=\frac{V_{qmax}Qn_{C}}{(\frac{V_{qmax}}{{\alpha\alpha}_{1}\phi_{1Pmax}}+Q)(4n_{C}-4n_{L}+3)}$	(S18)

$y=\frac{8\Gamma^{*}n_{C}-8\Gamma^{*}n_{L}+7\Gamma^{*}}{4n_{C}-4n_{L}+3}$	(S19)

These equations are valid for the case where mesophyll conductance is considered to be infinite, i.e., *C*_i_ = *C*_c_. Other equations are available to estimate the chloroplastic CO_2_ concentration while considering a finite mesophyll conductance (*62*).

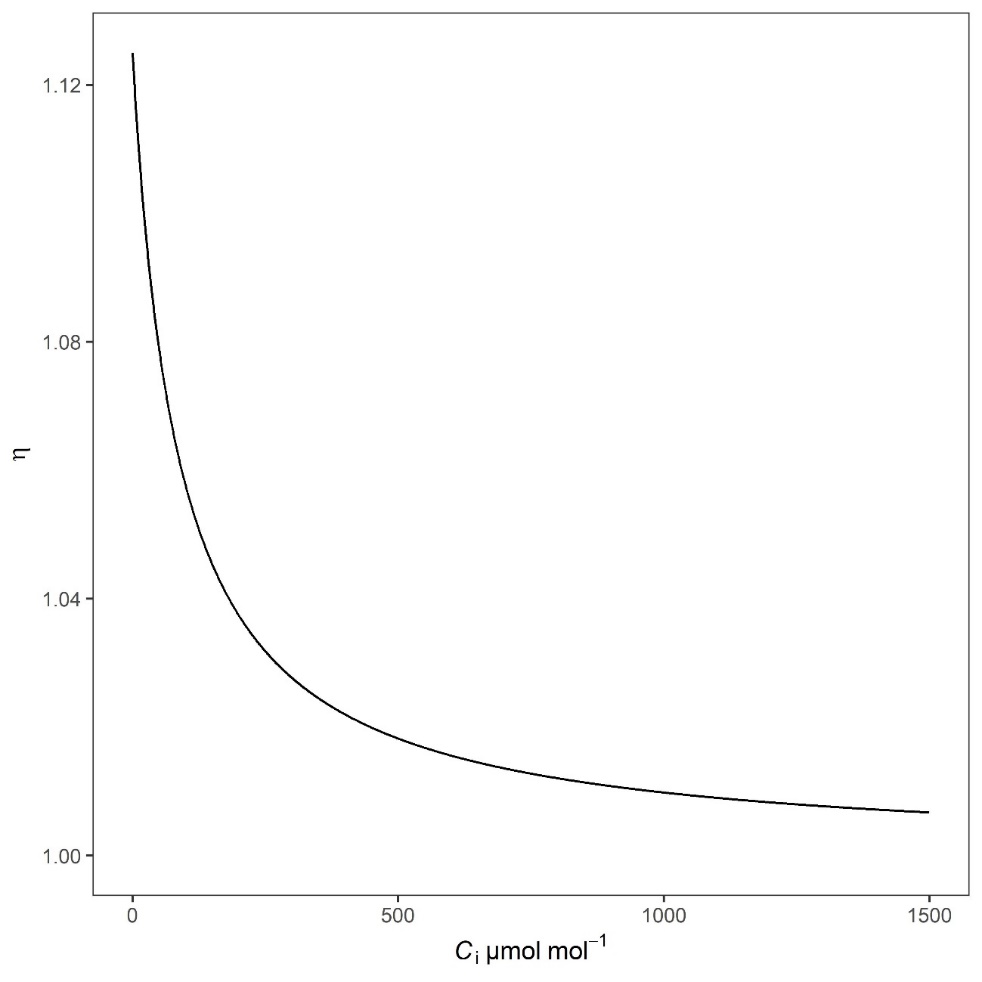

Fig. S1. Effect of the intercellular CO_2_ concentration (*C*_i_) on the variable *η* of the JB model.

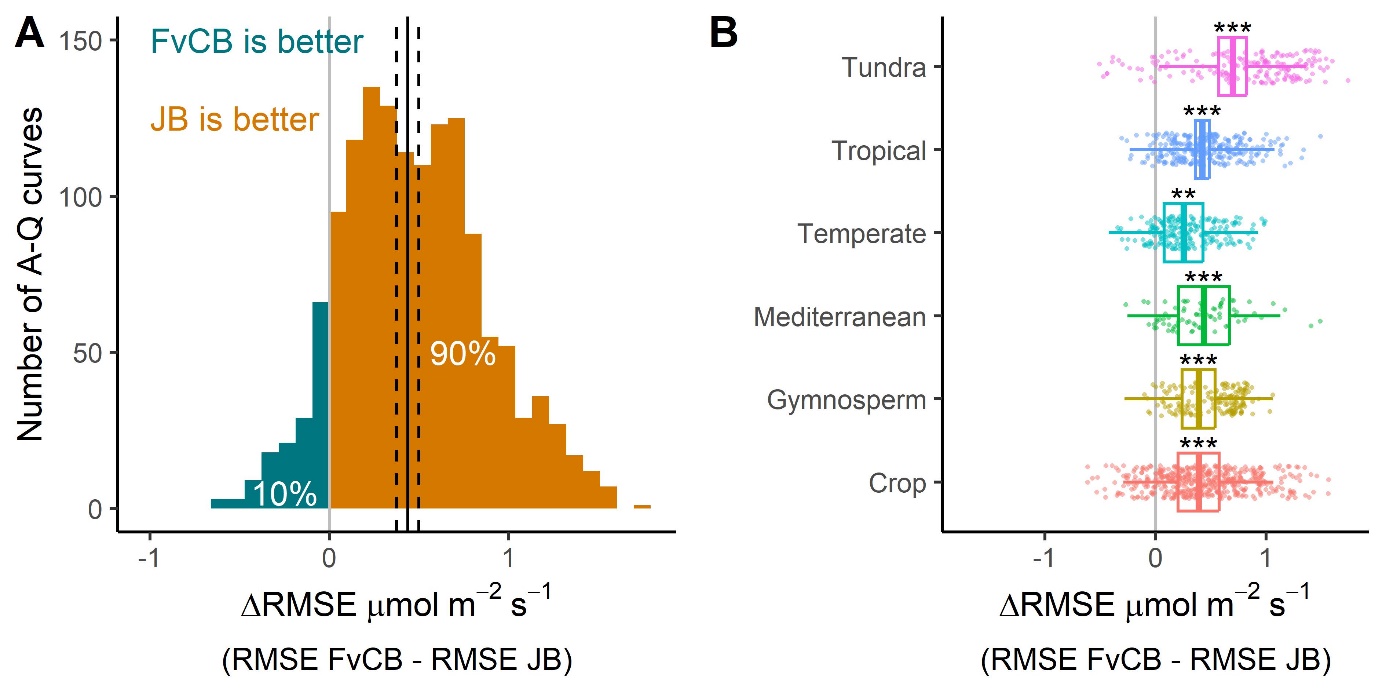
Fig. S2. Relative performance of the JB and FvCB models for representing the response of photosynthesis to irradiance (*A-Q* curves) in C3 species.

This figure is similar to Fig. 2 in the main text but with a different parameterization of the quantum yield ($\phi$ = 0.37) and curvature factor (*θ* = 0.9). The ΔRMSE represents the difference between the Root Mean Square Error between the FvCB and JB models, where a positive ΔRMSE indicates a lower error for the JB model and vice versa. **(A)** ΔRMSE distribution in the 1422 *A-Q* curves used in this analysis. The vertical black line represents the mean ΔRMSE calculated by a mixed model that considered the species and plant functional type random effects, and the dashed lines represent the standard error of the mixed model. **(B)** Relative performance of the FvCB and JB models for each plant functional type. The top and bottom of the boxes represent the 95% confidence interval and the line is the mean estimated by a mixed model with the species represented as a random effect. The whiskers show the 95% prediction interval. The asterisks show significant difference from 0 (* P<0.05, ** P<0.01, and *** P<0.001).

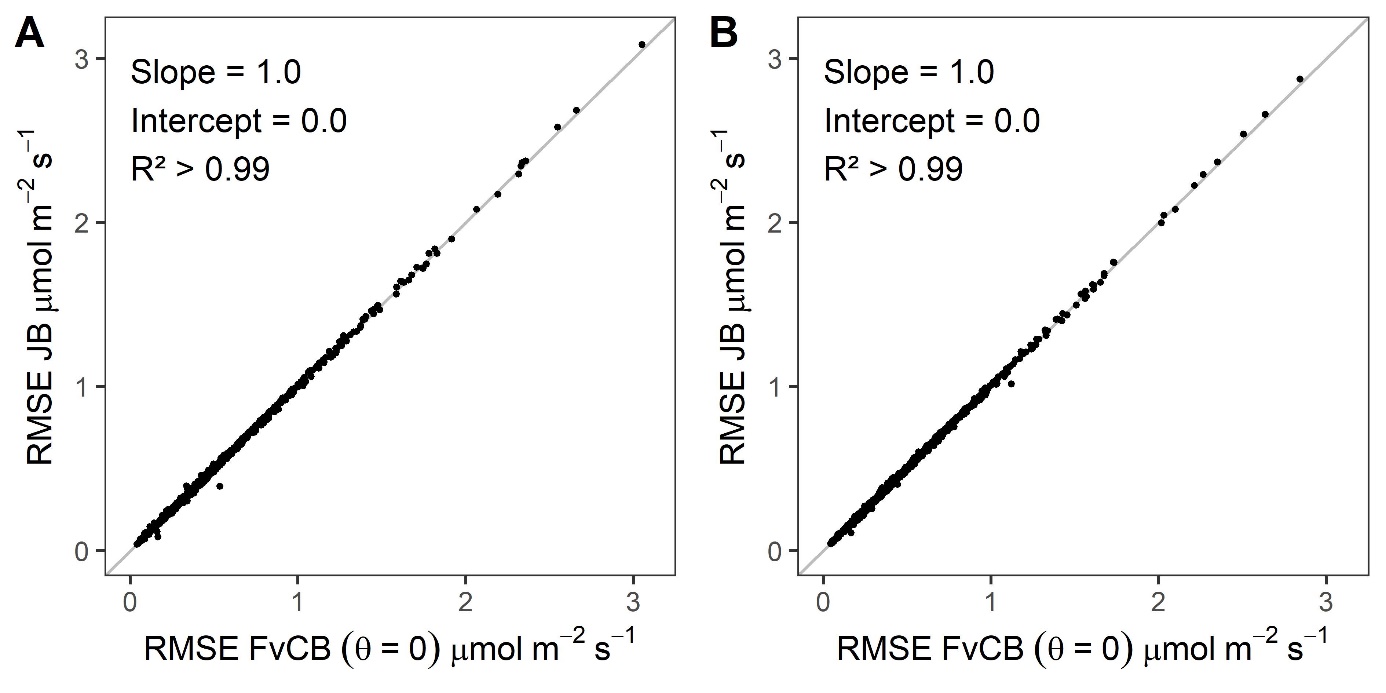
 Fig. S3. Comparison of model fits between the Johnson and Berry (JB) and Farquhar, von Caemmerer and Berry (FvCB) photosynthesis models.

Relationship between the root mean square error (RMSE) of the 1422 *A-Q* curves fitted with the JB and FvCB models, where the curvature factor (*θ*) in the FvCB model was fixed at 0. **(A)** Both models were parameterized with a quantum yield ($\phi$) of 0.42. **(B)** Both models were parameterized with $\phi$ = 0.37.

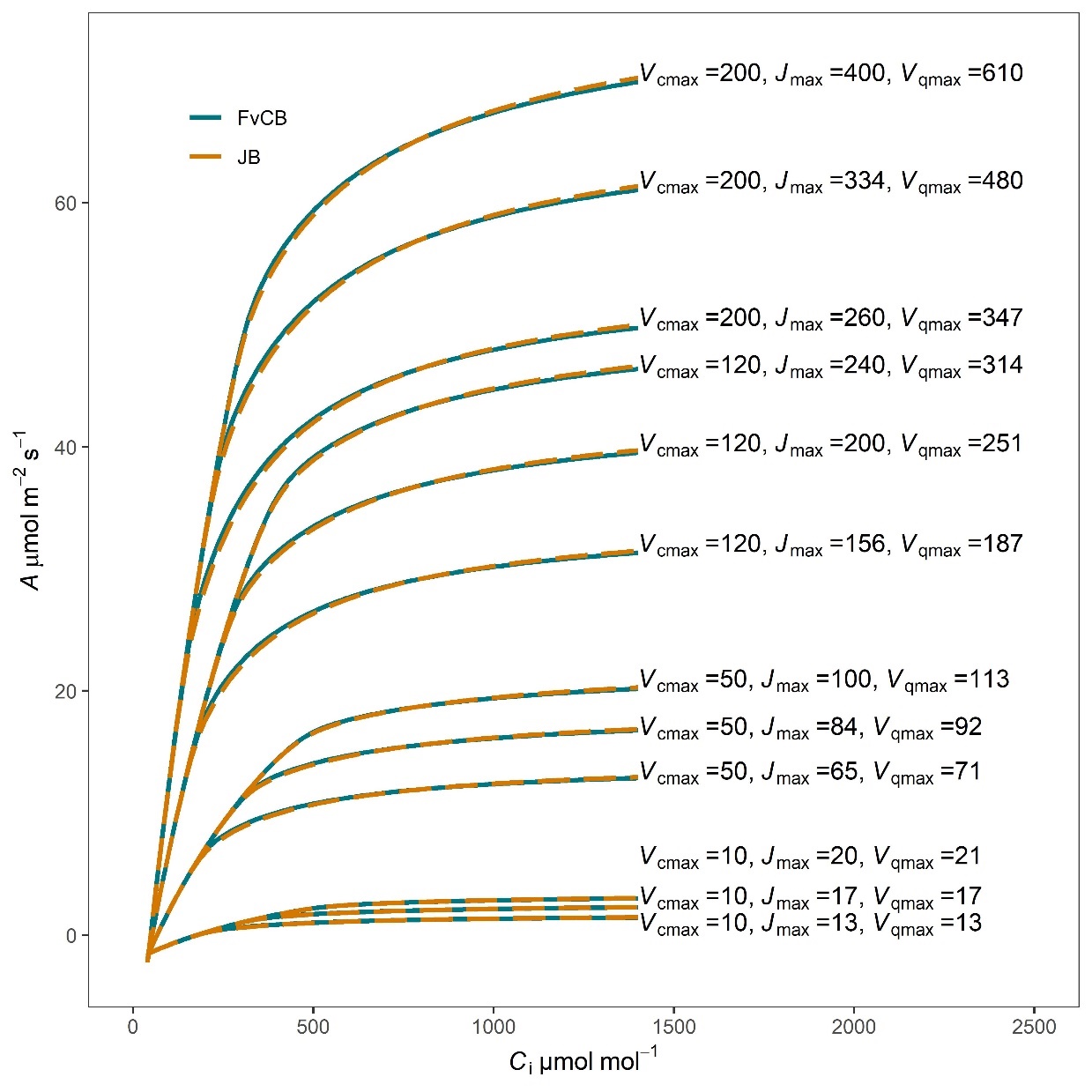

Fig. S4. Comparison of the response of photosynthesis to intercellular CO_2_ concentration (*A*-*C*_i_ curves) simulated with the FvCB and JB models of electron transport (*J*).

The maximum carboxylation rate of rubisco (*V*_cmax_) and the maximum electron transport rate (*J*_max_) were set at the values displayed in the panel (µmol m^-2^ s^-1^) for simulating each *A*-*C*_i_ curve with the FvCB model. The JB model parameters (α_1_ and *Cb*_6_*f*_max_) were adjusted to the FvCB model parameters using Equations (10) and (11).

Table S1. Terrestrial biosphere model (TBM) parameterization associated with the response of photosynthetic electron transport rate (*J*) to irradiance.

| TBM | *J* equation | *αΦ* | *Φ* | θ | Reference |
| --- | --- | --- | --- | --- | --- |
| - | Nonrectangular (Eqn S1) | 0.31 | 0.37 | 0.9 | Medlyn *et al.,* 2002 (*63*) |
| - | Nonrectangular (Eqn S1) |  | 0.425 | 0.7 | von Caemmerer *et al.,* 2009 (*48*) |
| - | Nonrectangular (Eqn S1) |  | 0.425 | 0.7 | Yin & Struik, 2009 (*64*) |
| FATES | Nonrectangular (Eqn S1) |  | 0.425 | 0.7 | Fisher *et al.,* 2015 (*17*) |
| CLM5 | Nonrectangular (Eqn S1) |  | 0.425 | 0.7 | Lawrence *et al.,* 2019 (*65*) |
| ED2 | Nonrectangular (Eqn S1) |  | 0.425 | 0.7 | Longo *et al.,* 2019 (*66*) |
| Orchidee | Nonrectangular (Eqn S1) |  | 0.425 | 0.7 | Krinner *et al.,* 2005 (*67*) |
| CABLE | Nonrectangular (Eqn S1) | 0.28 |  | 0.85 | Haverd *et al.,* 2018 (*68*) |
| JULES | Linear (Eqn S3) |  | 0.32 |  | Clark *et al.,* 2011 (*35*) |
| CTEM | Linear (Eqn S3) |  | 0.32 |  | Melton & Arora, 2016 (*69*) |
| JSBACH | Harley *et al.*(1992) (Eqn S2) | 0.28 |  |  | Mäkelä *et al.,* 2019 (*34*) |
| MedFATE | Nonrectangular (Eqn S1) | 0.30 |  | 0.9 | De Cáceres *et al.,* 2023 (*70*) |

Table S2. List of datasets used to compare the performance of the FvCB and JB models in species of the C3 photosynthetic pathway.

| Reference | PFT | N Species | N *A-Q* curves | Location | Environment | Method *A*-*Q* | N*A*-*C*_i_ curves | Irradiance *A*-*C*_i_ |
| --- | --- | --- | --- | --- | --- | --- | --- | --- |
| Burnett *et al.* 2019 (*28*) | Crop | 4 | 6 | USA, New York | Greenhouse | Rapid | - | - |
| Davidson *et al.* 2022 (*71*) | Temperate | 1 | 32 | USA, New York | Managed ecosystem | Slow | - | - |
| Davidson *et al.* 2023 (*72*) | Temperate | 7 | 185 | USA, New York | Forest | Slow | - | - |
| Davidson *et al.* 2024 (*73*) | Gymnosperm | 2 | 30 | USA, Alabama | Forest | Rapid | 35 | 1800 |
| Ely *et al.* 2024  (*74*) | Arctic | 11 | 62 | USA, Alaska | Tundra | Rapid | 68 | 1700 - 1800 |
| Fang *et al.* 2023  (*75*) | Crop | 1 | 242 | Netherlands | Greenhouse | Rapid | 234 | 1000 |
| Lamour *et al.* 2021  (*76*) | Tropical | 46 | 120 | Panama | Forest | Rapid | 112 | 400 - 2000 |
| Lamour *et al.* 2023  (*77*) | Tropical | 33 | 73 | Panama | Forest | Rapid | - | - |
| Lamour *et al.* 2024  (*78*) | Tropical | 26 | 82 | Brazil | Forest | Rapid | 31 | 1800 |
| Niinemets *et al.* 1999 (*79*) | Mediterranean | 1 | 20 | Portugal | Greenhouse | Rapid | - | - |
| Niinemets *et al.* 2004 (*80*) | Mediterranean | 3 | 56 | Portugal | Forest | Rapid | - | - |
| Niinemets *et al.* 2015 (*81*) | Crop | 1 | 8 | Estonia | Growth chamber | Rapid | - | - |
| Perez *et al.* 2024  (*82*) | Crop | 1 | 86 | France | Greenhouse | Rapid | - | - |
| Rogers *et al.* 2019 (*83*) | Arctic | 6 | 125 | USA, Alaska | Tundra | Rapid | 73 | 1800 - 2000 |
| Schmiege *et al.* 2021  (*84*) | Gymnosperm | 7 | 65 | Vietnam | Forest | Rapid | 48 | 400 - 1300 |
| Schmiege *et al.* 2023  (*85*) | Gymnosperm | 1 | 75 | USA, Alaska, New York | Forest | Rapid | - | - |
| Sun *et al.* 2012  (*86*) | Crop | 1 | 43 | Estonia | Growth chamber | Rapid | - | - |
| Vezy *et al.* 2024  (*87*) | Crop | 1 | 55 | France | Greenhouse | Rapid | - | - |
| Zhou *et al*. 2023*  (*88*) | Crop | 1 | 57 | Netherlands | Greenhouse | Rapid | - | - |

PFT: Plant Functional Type. N number of species or response curves included in this study after exclusion of some measurements detailed in the Materials and Methods. Method *A-Q*: type of protocol used to measure the light curves (*A-Q*): “Slow” corresponds to protocols with sufficient time for the stomata to acclimate to the light irradiance; “Rapid” corresponds to protocols where acclimation of stomata was not a requirement. Irradiance *A-C_i_*: irradiance levels at which the *A-C*_i_ curves were measured. * For Zhou et *al.* (2023) dataset, we only considered the non-modified genotypes
